# A Bayesian sequential learning framework to parameterise continuum models of melanoma invasion into human skin

**DOI:** 10.1101/284612

**Authors:** Alexander P Browning, Parvathi Haridas, Matthew J Simpson

## Abstract

We present a novel framework to parameterise a mathematical model of cell invasion that describes how a population of melanoma cells invades into human skin tissue. Using simple experimental data extracted from complex experimental images, we estimate three model parameters: (i) the melanoma cell proliferation rate, *λ*; (ii) the melanoma cell diffusivity, *D*; and (iii) *δ*, a constant that determines the rate that melanoma cells degrade the skin tissue. The Bayesian sequential learning framework involves a sequence of increasingly-sophisticated experimental data from: (i) a spatially uniform cell proliferation assay; (ii) a two-dimensional circular barrier assay; and, (iii) a three-dimensional invasion assay. The Bayesian sequential learning approach leads to well-defined parameter estimates. In contrast, taking a naive approach that attempts to estimate all parameters from a single set of images from the same experiment fails to produce meaningful results. Overall our approach to inference is simple-to-implement, computationally efficient, and well-suited for many cell biology phenomena that can be described by low dimensional continuum models using ordinary differential equations and partial differential equations. We anticipate that this Bayesian sequential learning framework will be relevant in other biological contexts where it is challenging to extract detailed, quantitative biological measurements from experimental images and so we must rely on using relatively simple measurements from complex images.

## 1 Introduction

Mathematical models of cell invasion may be expressed as coupled systems of partial differential equations where one component describes the density of invading cells, and another component describes the density of the receding tissues (Perumpanani et al., 1999; Gatenby et al.,1996; Landman et al., 1998; Smallbone et al., 2005; Anderson et al., 2008; Fasano et al., 2009; Swanson et al., 2011; Massey et al., 2012). Typically, models of cell invasion involve a population of motile, proliferative cells that release chemical signals to locally degrade surrounding tissues. These models have been applied to study malignant invasion (Gatenby et al., 1996) and developmental processes (Landman et al., 1998). While mathematical analysis of these models is relatively well established (e.g. Perumpanani et al., 1999), there are no standardised statistical protocols to parameterise these models using data from experimental images.

We consider the invasion of a population of metastatic melanoma cells into human skin tissue. Experimental images show that melanoma cells simultaneously migrate, proliferate and degrade the skin (Haridas et al., 2017b). To parameterise a parsimonious model of cell invasion we aim to infer three parameters: (i) the melanoma cell proliferation rate, *λ* > 0 [/h]; (ii) the melanoma cell diffusivity, *D* > 0 [*μ*m^2^/h]; and (iii) the rate at which melanoma cells degrade the tissue, *δ* > 0 [/h]. We take a likelihood-based Bayesian approach and work with a sequence of increasingly sophisticated experiments to identify these parameters. These experiments, listed here in order of increasing sophistication include: (i) a translationally-invariant proliferation assay; (ii) a barrier assay involving the spatial expansion of populations of melanoma cells; and (iii) the full invasion assay where proliferative and migratory melanoma cells degrade human skin tissues and then invade into the space created by receding tissues. A key outcome is to show that we obtain meaningful parameter estimates by working with relatively simple measurements from experimental images from sequence of increasingly-sophisticated experiments. In fact, we also show that naively attempting to identify *D, λ* and *δ* using only data from the most sophisticated, invasion experiment, leads to poorly-defined posterior distributions. While other studies have used information sequentially to help estimate parameters in mathematical models of cancer progression (Jain et al. 2014), we believe that this is the first study to use a Bayesian sequential learning approach to parameterise a mathematical model of cellular-invasion using spatial data extracted from experimental images.

One of the fundamental assumptions we make is that the melanoma cells are subject to the same individual-level mechanisms in all three experiments. For example, we make a standard assumption that melanoma cells undergo random migration and carrying capacity-limited proliferation in all three experiments. This does not mean that we observe the exact same population-level behaviour in all three experiments. For example, we see larger net growth of the melanoma population in the proliferation assay and barrier assay compared to the invasion assay. This is because melanoma cells in the invasion assay must compete for space with both the human skin tissue and other melanoma cells whereas in the barrier and proliferation assays there is no skin tissue and the only competition for space is between the melanoma cells.

## 2 Experimental methods

All experiments use the SK-MEL-28 metastatic human melanoma cell line (Carey et al., 1976). All experimental data are summarised in the Supporting Material.

### 2.1 Type 1: Proliferation assay

A proliferation assay involves uniformly placing a population of cells, at low density, on a two-dimensional substrate. Cells migrate and proliferate, and the density of the monolayer increases (Browning et al., 2017). On average, proliferation assays are translationally invariant since the population of cells is distributed uniformly. Therefore, we simply count the number of cells in the field of view to characterise the increase in population density over time. We use images and data from Haridas et al. (2017a). The population growth is quantified by counting the number of cells in several regions, and dividing by the area of the region and the carrying capacity density, *K* = 2.8 × 10^−3^ cells/*μ*m^2^ (Haridas et al., 2017a), to give an estimate of the nondimensional cell density at *t* = 0,24 and 48 h. We consider three identically-prepared experimental replicates (Haridas et al., 2017a), and 26 subregions per replicate to give 78 nondimensional density estimates per time point. Images in Figure 1(a)-(c) show the growth process, and data in Figure 1(d) summarises the data.

**Fig. 1.**
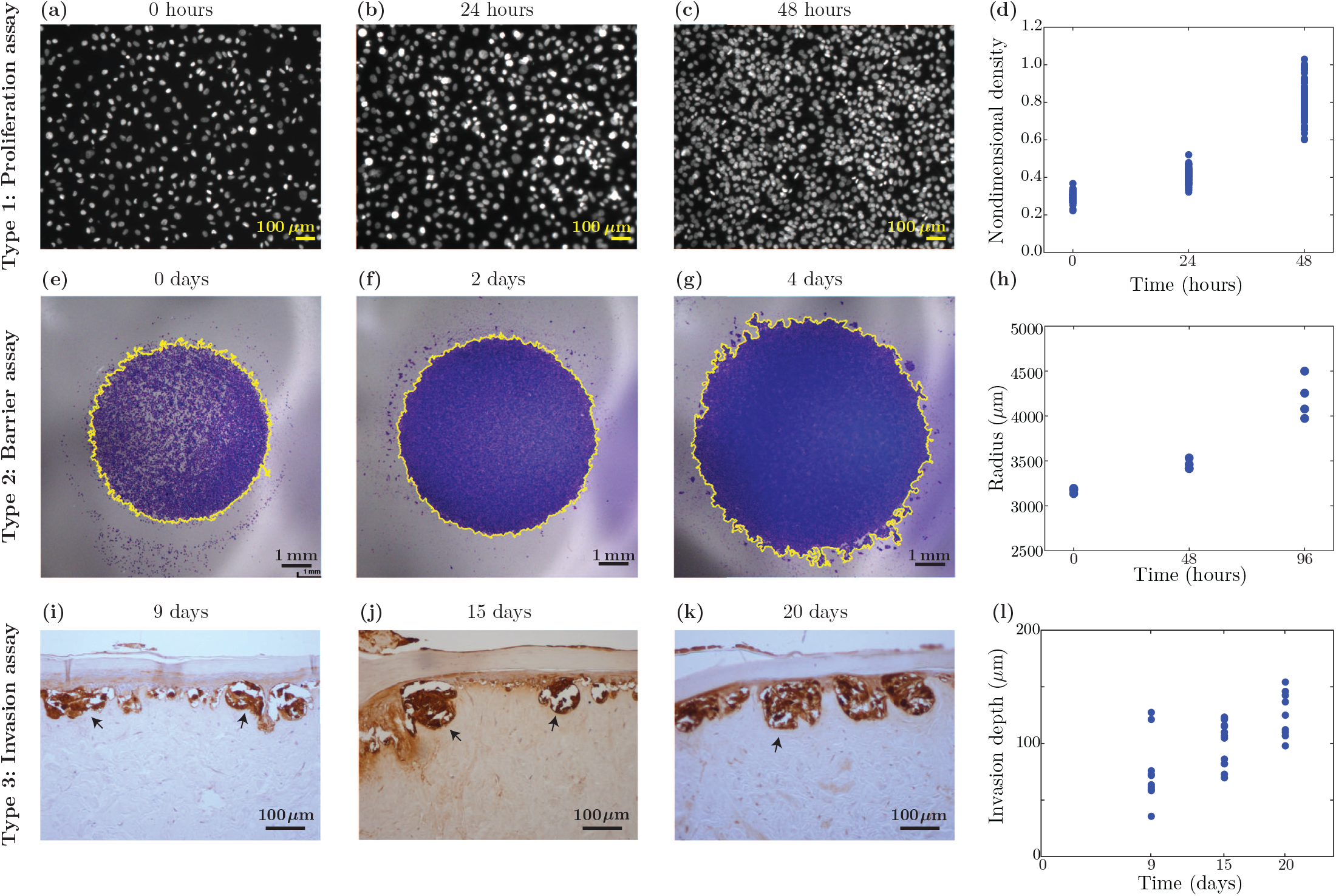
Experimental images and data summary. (a)-(c) Typical images from the proliferation assay with SK-MEL-28 cells. The nondimensional density increases with time, as summarised in (d). (e)-(g) Typical images from the circular barrier assay with SK-MEL-28 cells, with data in (h) showing the increase in radius of the spreading circular population. (i)-(k) Typical images from the invasion assay with SK-MEL-28 cells, with data in (l) showing the increase in depth of invasion. Images in (a)-(c) are reproduced from Haridas et al. (2017a); images in (e)-(g) are reproduced from Haridas et al. (2018); and images in (i)-(k) are reproduced from Haridas et al. (2017b). The differences in colour and appearance between the experimental images are due to differences in staining techniques between the three different types of experiments.

### 2.2 Type 2: Barrier assay

A circular barrier assay is initiated by uniformly placing a population of cells inside a circular barrier (Treloar et al., 2013a). The barrier is lifted and the population of cells spreads outwards across a two-dimensional surface. Figure 1(e) shows that the initial population of 20,000 melanoma cells is confined to a circular region with a diameter of approximately 6 mm. This means that the initial density of the monolayer inside the barrier is 20,000/(*π*3000^2^) ≈ 7.07 × 10^−3^ cells/*μ*m^2^, corresponding to an initial nondimensional density of 20,000/(*Kπ*3000^2^), giving approximately 0.25. Over four days the population spreads to occupy a circular region with a diameter of approximately 9 mm (Haridas et al., 2018). The key difference between the circular barrier assay and the proliferation assay is that the proliferation assay is translationally invariant whereas the barrier assay is not, as the barrier assay involves moving fronts of cells. Therefore, the proliferation assay is well-suited for estimating the cell proliferation rate, and the circular barrier assay is well-suited for estimating the cell diffusivity.

Automated image processing, implemented in MATLAB (Mathworks, 2018a), is used to quantify the spreading of the cell population by estimating the position of the leading edge. This involves applying steps 1-7 from Algorithm 1, which are shown in Figure 2(a)-(f) (Treloar et al., 2013b). Following this initial process, we obtain a mean pixel density profile as a function of the radius using the procedure outlined in steps 8-14 of Algorithm 1. This second series of steps are outlined visually in Figure 2(g)-(h). To summarise each experimental image, we first consider that the scaled mean pixel density at the centre of the assay is unity. We then estimate the position of the leading edge of the spreading population to be the radius at which the scaled mean pixel density of 1% of the initial maximum pixel density. This allows us to estimate the position of the leading edge of the spreading population where the density is approximately 1% of the maximum initial density. The threshold of 1% has been shown, in previous studies, to give a reliable measure of the extent of spatial spreading (Treloar et al., 2013b). This process is repeated for four identically-prepared barrier assays, at each time step, and the data is summarised in Figure 1(h).

#### Algorithm 1 Quantifying experimental images from a circular barrier assay using the image processing toolbox in MATLAB (Mathworks, 2018a)

1. Load and crop image using `imread()`.
2. Convert image to grayscale using `rgb2gray().`
3. Obtain the gradient mask using the Sobel method, edge(· ,‘Sobel’, *γ*thresh), where `thresh` is the `MATLAB` suggested threshold, and *γ* is an adjustment parameter. We initially fix *γ* = 1 for each image, and adjust as necessary.
4. Obtain dilated binary mask using `imdilate` and `strel` with a `‘disk’` structuring element.
5. Fill holes in the mask using `imfill(· ,‘holes’)`.
6. Smooth the mask using `imerode` and `strel` using a `‘disk’` shaped structuring element.
7. Clear border objects using `imclearborder` and remove small areas using `bwareaopen`.
8. Use `regionprops(·,’centroid’)` to obtain the coordinates of the centre of the area.
9. Determine the distance of each pixel in the region from the calculated centre.
10. Use `histogram()` to obtain the distribution of distances using *Δ* = 5 *μ*m.
11. Scale by the area of each ‘bin’, 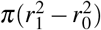, where *r*_0_ and *r*_1_ are the radius edges of each bin.
12. Scale using the length scale of each image to determine the density profile.
13. Smooth so that the density at small radii is 1.
14. Obtain the profile at all distances as required using `interpl()` and the `‘spline’` method.

**Fig. 2.**
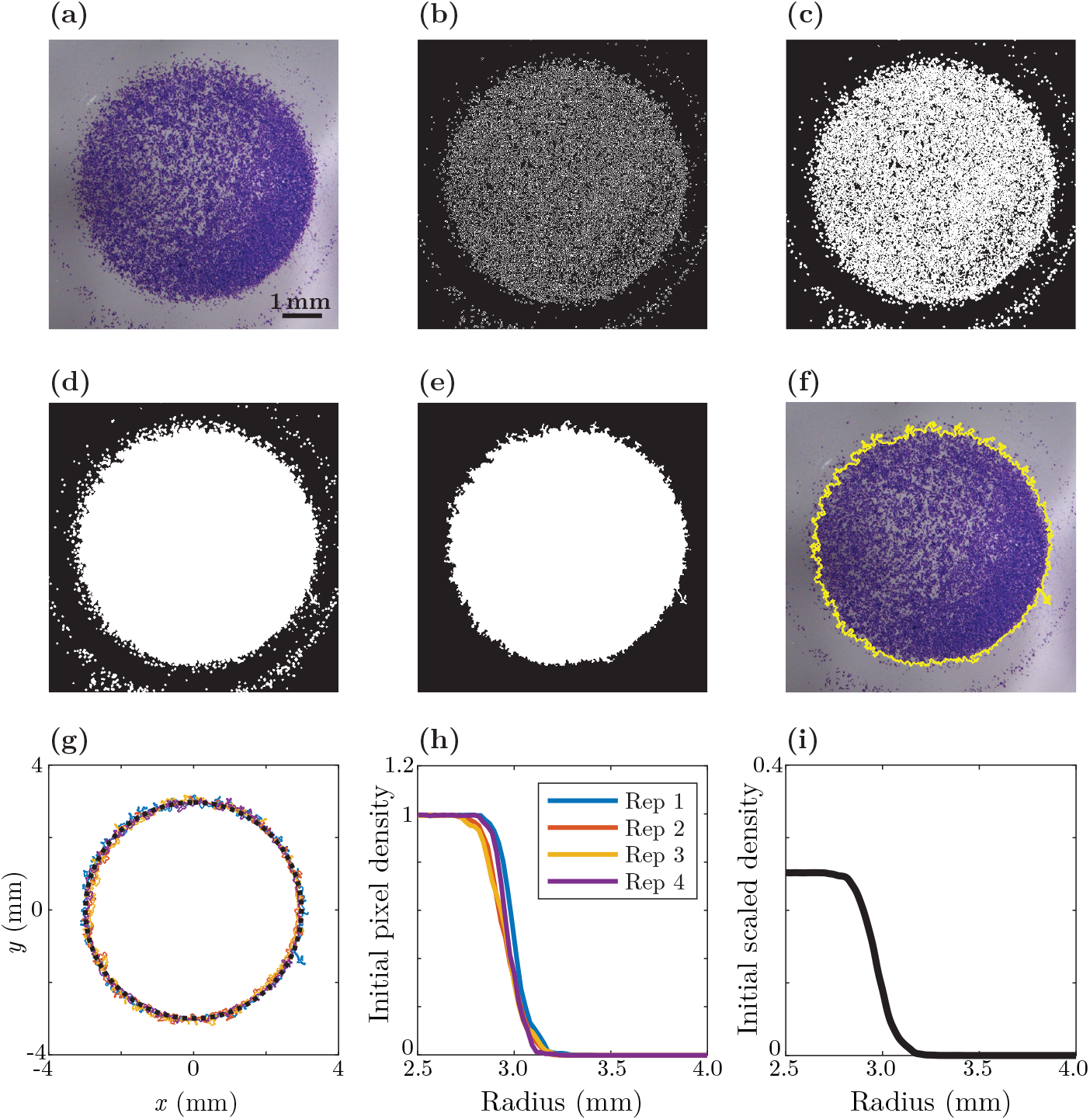
Automated image processing for the circular barrier assay. (a)-(f) Stages in the automatic leading edge detection algorithm: (a) original image; (b) grayscale gradient mask; (c) dilated binary mask; (d) binary image with holes filled; (e) clear binary image; (f) original image with detected leading edge superimposed, showing a good match. (g)–(l) Stages in determining the location of the leading edge using several identically-prepared images: (g) leading edge for each experimental replicate at *t* = 0 h, with the centre corresponding to the centre of mass of each region; (h) average pixel density profile as a function of radius, *r*, for each experimental replicate; (i) average pixel density, scaled relative to the initial density of 20,000 cells within a circle of diameter 6 mm, to represent the scaled pixel density as a function of *r*.

### 2.3 Type 3: Invasion assay

The invasion assay involves observing how a monolayer of melanoma cells invade into human de-epidermised dermis prepared from discarded human skin tissue, as described by Haridas et al. (2017b). Primary skin cells are used in the invasion assay to ensure the formation of a stratified epidermis with a basement membrane (Haridas et al., 2017b). Vertical invasion of melanoma cells, downward through the basement membrane into the dermis, is observed. The depth of invasion beyond the basement membrane is estimated using immunohistochemistry. Measurements quantify the depth of invasion into the dermis after 9, 15 and 20 days, thereby providing temporal information about the invasion process. The experimental protocol of Haridas et al. (2017b) involves labelling all melanoma cells, and it is difficult to distinguish between closely located individual melanoma cells within the complex skin environment. Therefore, the simplest measurement to characterise these experiments is to record the depth of the deepest positively-stained melanoma cell in each image. This measurement is both mathematically and biologically useful. Firstly, from a mathematical point of view, the continuum model we use gives rise to a sharp-fronted solution meaning that the maximum depth of invasion is a convenient way to connect the experimental data to the solution of the mathematical model. Secondly, from a biological point of view, the maximum depth of invasion is known clinically important (Haridas et al. 2017b). A summary of images in Figure 1(i)-(j) show the melanoma cells invading into the skin tissue. These images show that the invasion of the melanoma cell population is closely associated with the receding skin tissues. Data in Figure 1(l) shows multiple measurements of the maximum depth of invasion from many identically prepared experimental replicates using different human de-epidermised dermis. Here the experiments are summarised by measuring the distance between the deepest melanoma cells and the boundary between the epidermis and the dermis (Haridas et al., 2017b). Nine identically prepared invasion experi ments are performed for each time point, and the distribution of invasion depths are presented in Figure 1(l).

## 3 Mathematical models

A detailed discussion about the development of the mathematical models, the key assumptions underlying these models, and their nondimensionalisation, is given in the Supporting Material. In brief, we use a sequence of related models that we present here in order of increasing sophistication. In all cases, we choose nondimensionalise the dependent variables, but work with dimensional independent variables and dimensional parameters.

### 3.1 Model 1: Temporal one-component model for the proliferation assay

For a spatially uniform population of melanoma cells in the absence of skin tissue, we make the standard assumption that the population grows logistically (Sengers et al., 2007; Maini et al., 2004; Swanson et al., 2011).

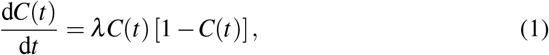

so that

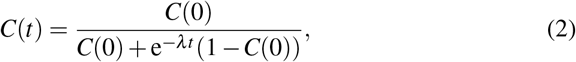

where *C*(*t*) is the nondimensional density of the monolayer at time *t*.

### 3.2 Model 2: Spatial and temporal one-component model for the barrier assay

We assume that a population of motile and proliferative melanoma cells spreads according to the Fisher-Kolmogorov model (Sengers et al., 2007; Maini et al., 2004; Swanson et al., 2011), provided that there is no skin tissue present. Written in radial coordinates, we have

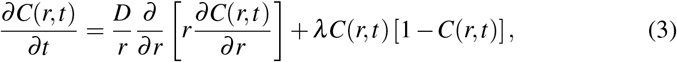

where *r* > 0 is the radial position.

### 3.3 Model 3: Spatial and temporal two-component model for the invasion assay

The full mathematical model of the invasion assay is given by,

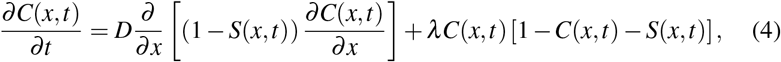

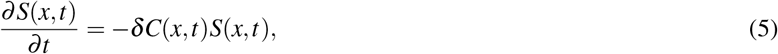

where *C*(*x, t*) is the nondimensional density of melanoma cells, *S*(*x, t*) is the nondimensional density of skin tissue and *x* is the vertical depth into the tissue. In brief, the movement of the melanoma cells is governed by a nonlinear diffusion term where the nonlinear diffusivity is a decreasing function of the skin density so that melanoma cells are unable to diffuse when the skin is at maximum density. The proliferation of melanoma cells is logistic, and crowding effects are incorporated so that when the total density of skin and melanoma cells are at maximum density the net proliferation rate is zero. The skin tissues degrade when in contact with melanoma cells. Other choices for the form of the nonlinear diffusivity function and the sigmoid proliferation model are possible, and we briefly discuss these options in the Conclusions.

The three models that we consider are closely related. To see this, setting *S*(*x, t*) = 0 in Equations (4)-(5) leads to the Fisher-Kolmogorov equation which, when written in radial coordinates, gives Equation (3). Similarly, setting *S*(*x, t*) = 0 and *∂C*(*x, t*)/*∂x* = 0 in Equations (4)-(5) leads to Equation (1). The methods used to solve Equations (3)–(5) are outlined in the Supporting Material.

A notable feature of the mathematical models is that we choose to deal with dimensional independent variables and non-dimensional dependent variables. This is a deliberate choice that we find to be both practical and insightful. For example, when we work with experimental images it is straightforward to measure the relevant physical dimensions, such as the relevant length scales and relevant time scales. For example, the images in Figure 1(e)-(g) show that the fronts of cells move approximately 1000-2000 *μ*m over a duration of four days. Therefore, it is most biologically meaningful for us to deal with dimensional time with units of [h], dimensional position with units of [*μ*m], dimensional diffusivities with units of [*μ*m^2^/h] and dimensional proliferation rates with units of [/h]. Our interpretation of the dependent variable, *C*(*x, t*), however, is very different. Typically, in continuum models of cell migration and cell invasion, the dependent variable is the cell density which is bounded by some maximum carrying capacity density (e.g. Maini et al. 2004a; Maini et al. 2004b). It is well-known that extracting estimates dimensional cell densities as a function of position and time from experiments can be extremely challenging since it often replies on manually counting cells to construct cell density profiles (Treloar et al. 2013; Vittadello et al. 2018). We believe that an attractive feature of our approach is that we do not rely on counting individual cells to construct cell density profiles for the invasion assay or the barrier assay. It is convenient to avoid counting individual cells because this procedure is both extremely time consuming and technically challenging when dealing with high-density experiments. Therefore, to make our procedure as useful as possible to others, we rely only on dealing with leading edge data because this data is far easier and faster to compute (Treloar et al. 2013). The simplest way for us to interpret the density information in the invasion assay and the barrier assay using the mathematical models is to non-dimensionalise the dependent variable in the mathematical models and treat the position of the leading edge data as corresponding to the low density edge of the cell density profile where *C*(*x, t*) ≪ 1. Therefore, while it might at first appear to be unusual to deal with dimensional independent variables and dimensionless dependent variables, this choice is driven purely by biological and practical reasons which we believe to hold across of range of cell biology problems where the aim is to match the solution of a mathematical model to mimic complicated details in a biological image.

Another feature of our mathematical models is that the three models deal with different spatial dimensions. For example, the proliferation assay is constructed so that the measurements of cell density are translationally invariant, and it is standard to extract density information from these kinds of experiments and to treat that information as being dependent on time only (e.g. Cai et al. 2007; Tremel et al. 2009). In the barrier assay, the experiments are designed so that we observe the movement of cell fronts and so the experiments are not translationally invariant since density depends upon position. In this situation it is standard to report the density of cells as a function of position in the two-dimensional plane, (*x, y*), or as a function of the radial coordinate, *r*^2^ = *x*^2^ + *y*^2^, for radially symmetric problems (Treloar et al. 2013a; Treloar et al. 2013b). The invasion assay is clearly three-dimensional since cells move both horizontally and vertically in the human skin tissues (Haridas et al. 2017b; Haridas et al. 2018). Therefore, in general, one could use a three-dimensional Cartesian coordinate system to describe the cell density as a function of position. However, the geometry of the experiments allows us to simplify this description to describe the cell density as a function of the vertical coordinate only. In the experiments melanoma cells are uniformly placed onto the surface of the skin tissues in a circular barrier of diameter 6000 *μ*m (Haridas et al. 2017b; Haridas et al. 2018). Measurements of the depth of invasion are intentionally made under the centre of the circular area, and we find that the depth of invasion is relatively small compared to the horizontal extent of the barrier. In summary, cells invade vertically into the skin approximately 150 *μ*m over 20 days. This means that the vertical length scale is much smaller than the horizontal length scale since 150/6000 = 0.025 ≪ 1. Under these situations we can reduce the description of a three-dimensional transport process to a one-dimensional coordinate system based on a geometric argument (Simpson et al. 2017). We acknowledge that had the measurements of invasion been made towards the edge of the 6000 *μ*m radius of the barrier, or had the invasion process taken place over larger vertical length scales, it would be appropriate to use a two- or three-dimensional model to describe this situation (Simpson et al. 2017).

### 3.4 Initial conditions

For each model we specify an initial condition to match the initial experimental measurements.

> **Model 1**. *C*(0) is the average nondimensional density measured at *t* = 0.
>
> **Model 2**. Algorithm 1 gives the scaled pixel density profile as a function of *r*, for images at *t* = 0. With this information we compute the average the density profile across all experimental replicates, and re-scale so that the nondimensional density at the centre of the circular population corresponds to placing 20,000 cells of diameter 20 *μ*m into a circular barrier of radius 3000 *μ*m.
>
> **Model 3**. The invasion into the dermis commences approximately 4 days after the invasion assay is initialised (Haridas et al., 2017b). To capture this we assume the density of a monolayer of melanoma cells initially placed onto the surface of the skin tissue grows logistically over the first 4 days, giving the initial nondimensional density of melanoma cells at the top of the tissue to be approximately 0.78. To model the spatial aspects of the invasion assay, we assume the monolayer of melanoma cells is 20 *μ*m thick (Haridas et al., 2017a), giving *C*(*x*, 0) = 0.78 for −20 < *x* < 0, and *C*(*x*, 0) = 0 for *x* > 0. We assume that the density of skin tissue is the maximum possible density, *S*(*x*, 0) = 0 for −20 < *x* < 0, and *S*(*x*, 0) = 1 for *x* > 0.

### 3.5 Boundary conditions

For the spatial models, model 2 and 3, we specify boundary conditions that are consistent with the experimental design.

> **Model 2**. The initial radius of the spreading population is *r* = 3000 *μ*m and we find that after four days the radius of the spreading population is no more than approximately 4500 *μ*m. Since the radius of the wells in the 24-well plate are approximately 7800 *μ*m, the leading edge of the spreading profile does not touch the edge of the well. To account for this we solve Equation (3) on 0 < *r* < *R* with *∂C*/*∂r* = 0 at *R* = 7800 *μ*m to account for the physical boundary at the edge of the circular well. We also set *∂C*/*∂r* = 0 at *r* = 0 to account for the radial symmetry.
>
> **Model 3**. The depth of the tissues in Figure 2 is several millimetres (not shown), yet the invasion depth of the melanoma cells is only 150 *μ*m so that the boundary condition at the bottom of the tissue does not affect the solution of Equations(4)-(5) on the time scale of these experiments. To account for this we solve the equation governing *C*(*x, t*) in Equation (4) on −20 < *x* < *L*, where we set *C*(*L, t*) = 0 for *L* = 1000 *μ*m and we note that our results are insensitive to this choice of *L*. We also note that Equation (5) involves no spatial derivative terms so there is no boundary conditions required to solve this equation.

### 3.6 Summarising model observations

To connect the models with the experimental measurements, we summarise key features of the model solutions that can be matched with simple, objective measurements from the experimental images. Our aim is to estimate ***Θ*** = 〈*λ, D, δ*〉. Throughout we denote ***M**_k_*(*t; **Θ***) as a summarised model observation from model *k*, at time *t*. Here, *k* = 1,2 or 3. For each model we summarise the observation as follows:

> **Model 1**. The density:
>
>
> 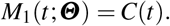
>
> **Model 2**. The radius of the leading edge:
>
>
> 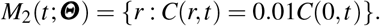
>
> **Model 3**. The depth of the front of melanoma cells:
>
>
> 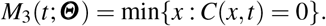

These data, which summarise the predictions of the model, are chosen because they are objective, simple measurements that can be obtained from experimental images.

## 4 Statistical inference

Taking a Bayesian approach we consider both the model parameters, ***Θ***, and experimental observations to be random variables (Gelman et al., 2004; Toni et al., 2009; Fearnhead et al., 2012; Collis et al., 2017; Browning et al., 2018; Daly et al., 2018). We consider that the deterministic models capture the *expected* behaviour, and that the experimental data in Figure 1(d),(h),(l) characterises some measurable experimental variability (Warne et al., 2017). Therefore, we make the natural assumption that the experimental observations are normally distributed about the solution of the corresponding model (Collis et al., 2017; Warne et al., 2017), and we assume the observation variance within each experiment is a constant, which we denote 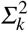.

Before we make any experimental observations, our knowledge about the parameters is contained within the prior distribution, *p*(***Θ***). We denote a sequence of experimental observations 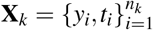, where *y_i_* is an experimental observation from model *k* at time *t_i_* and *n_k_* is the number of times that experimental data is recorded for experiment type *k*. We may therefore express the likelihood, 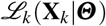, or probability density of the experimental data given the model parameters as

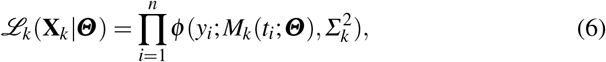

where ***M**_k_*(*t_i_; **Θ***) is a summary model observation at time *t_i_* from model *k*, and *ϕ* denotes a normal probability density with mean ***M**_k_*(*t_i_; **Θ***) and variance 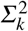. We approximate 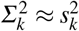, where 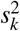 is a pooled sample variance of the time-grouped observations for each experimental data set. That is, we calculate the variance of the pooled sample for each experiment, after the mean of each time group has been subtracted. Specifically,

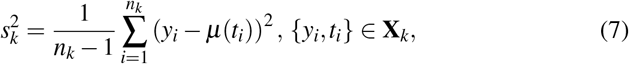

where *μ* (*t_i_*) is the mean of the set of experimental observations made at time *t_i_*. This assumption allows for a different mean between each group of data at different time points.

Using Bayes’ theorem, we update our knowledge of the parameters to form a posterior distribution,

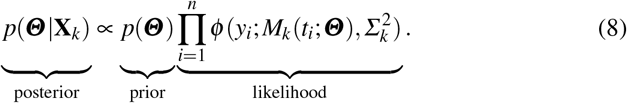

A key element of this study is to contrast how a posterior distribution using a uniform prior differs from an *informed* prior that is built sequentially. We consider a uniformly distributed prior defined over a sufficiently large parameter space so that all biologically feasible parameter combinations are covered. We do not specify the domain of the prior, and we use the scaled posterior distribution to obtain information such as maximum likelihood estimates.

When forming posterior distributions using *informed* prior distributions, *p_k_*(***Θ*** |**X**_*k*_), we use a sequential approach. That is, we specify the prior distribution for model *k* to be the posterior distribution for model *k* − 1,

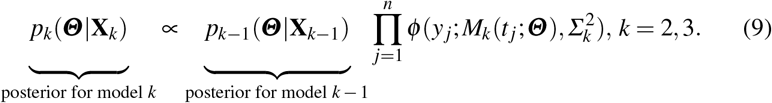

Mathematically, the posterior distribution formed for model *k* using this technique is equivalent to the posterior distribution given data up to experiment type *k*. That is,

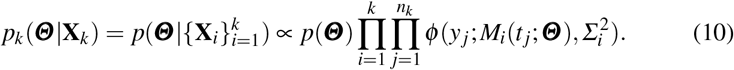

In practise it is simpler to apply Equation (9) to form these posterior distributions rather than Equation (10). For example, Model 1 only depends on *λ*, a single element of ***Θ***. Therefore, the other components of the posterior distribution, *D* and *δ*, remain uniform when we work with Equation (9) for model 1. It is relatively straightforward to find the posterior support for a single parameter rather than finding the posterior support for multiple parameters simultaneously. As more parameters are incorporated in successive models, in this case one at-a-time, the search for the posterior support remains a simple task. In contrast, and we as will demonstrate, it is both practically challenging and computationally inefficient to find the posterior support for Model 3 directly, since it depends on all three components of ***Θ***. As a result, our sequential approach allows us to estimate a three-dimensional posterior distribution easily and efficiently, whereas the direct approach fails to produce meaningful results.

When presenting posterior distributions, we calculate the posterior distribution exactly at points on a relatively coarse square discretisation of the parameter space (Supporting Material). Our choice of discretisation allows us to calculate maximum likelihood estimates accurately to two significant figures, without further refinement. We then use a spline interpolation (Mathworks, 2018b) to both enhance the resolution of the posterior distributions and to approximate the posterior density at points that do not lie on the square discretisation, as required.

### 4.1 Credible regions

To summarise the posterior distributions we compute and show credible regions. We first calculate the total scaled posterior distribution volume, 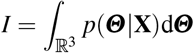, using quadrature, on the smoothed posterior distribution. In this work we use the rectangle rule to approximate the integrals. This procedure is relevant for the informed sequential posterior distributions since it is visually obvious that we have covered the support of the distribution. In contrast, this approach is not possible for the posterior distributions that use a uniform prior, since we have not calculated the posterior density through the entire support.

Figure 3(a)-(b) illustrates how we calculate the credible region *Q*, bounded by *q*, for one- and two-dimensional posterior distributions. The 1 − *α* credible region of *p*(***Θ***|**X**) is

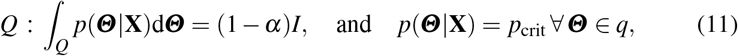

for *α* ∈ [0,1]. This means that the total posterior density within *Q* is 1 − *α*, with constant posterior density on the boundary, *p*_crit_. We approximate this region by estimating *p*_crit_ such that

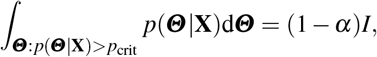

where the integral is estimated using quadrature, in this case the rectangle rule. For all results we set *α* = 0.05 to calculate a 95% credible region.

**Fig. 3.**
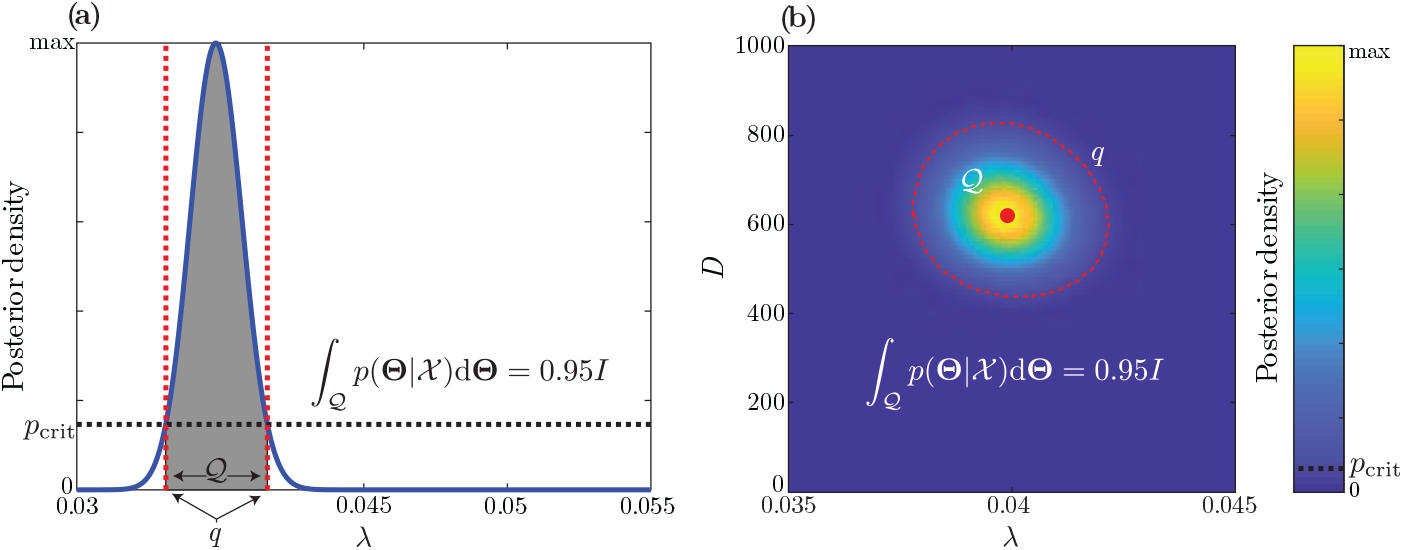
Example credible region calculations. In each case the boundary of the interval or region, denoted *q*, has constant posterior density. The total area under in the one-dimensional case, or total volume in the two-dimensional case is 0.95I. Extending the credible region calculation in (b) to deal with higher-dimensional posterior distributions is straightforward. The boundary of the 95% credible region is indicated with a dashed red line.

When calculating a credible region, we increase the accuracy of the reported credible interval by smoothing the probability density function using the `interp` function in MATLAB with a cubic spline interpolation (Mathworks, 2018b). This approach allows us to increase the accuracy of the credible interval estimates further than our relatively coarse discretisation of the parameter space would otherwise allow. The details of the discretisations are given in the Supporting Material document. In the Supporting Material document we show how this processing provides a similar, but visually smoother approximation to credible regions than what would otherwise be possible with the relatively coarse discretisation.

### 4.2 Prediction intervals

To demonstrate uncertainty in the model predictions, we approximate and present prediction intervals along with a model prediction produced using the mode of the posterior distribution. It should be noted that the borders of the prediction intervals we present are not model realisations. Rather, these intervals correspond to the interval containing 95% of model realisations. Prediction intervals are calculated by sampling 50,000 parameter combinations from the posterior distribution and solving the appropriate model for each combination. We continue our assumption that the model captures a normally-distributed experimental variability by adding Gaussian noise to each model realisation, independently at each time point. For each time point, we use the `ksdensity` function in MATLAB (Mathworks, 2018c) to form a probability density function, from which we follow our previously outlined procedure to approximate a credible interval.

## 5 Results and Discussion

The first step is to estimate *λ* from Equation (1). Results in Figure 4(a) show that we arrive at a well-defined, approximately symmetric posterior. The posterior mode is 0.040 /h and 95% credible interval 0.038 < *λ* < 0.042 /h. Our estimate of the mode corresponds to a doubling time of ln(2)/0.04 ≈ 17 h, which is fairly typical for a melanoma cell line (Treloar et al., 2013a). It is also useful to note that the posterior support for *λ* is relatively narrow. In our preliminary investigations (not shown), we originally explore the interval 0 < *λ* < 0.2 /h, but since we find non-zero posterior density for just a small region within this interval we present results in Figure 4(a) on just 0.03 < *λ* < 0.05 /h.

**Fig. 4.**
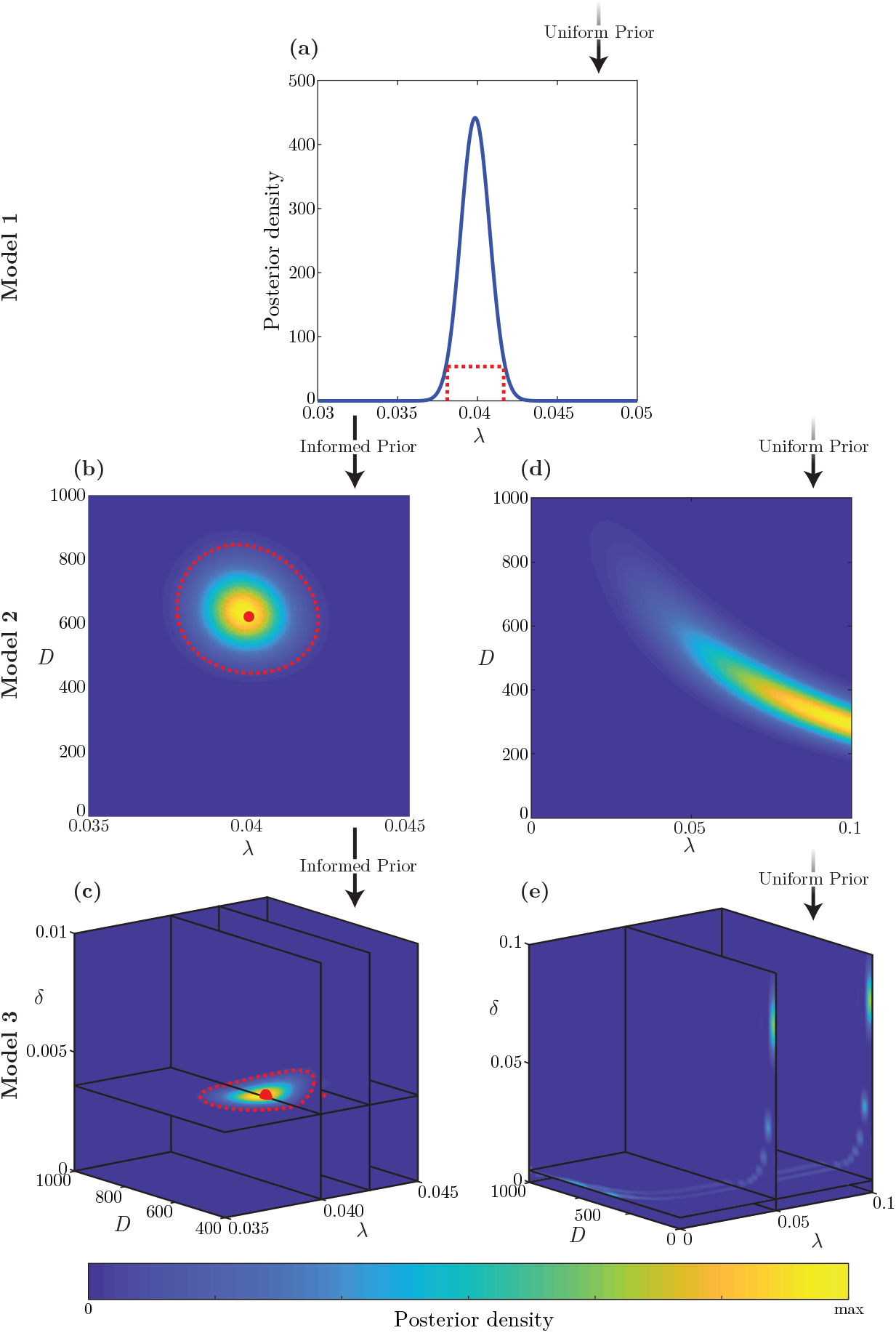
Posterior distributions. Posterior distributions produced for each model and each experiment: (a) model 1; (b),(d) model 2; and, (c), (e) model 3. (a), (d) and (e) Posterior distributions using a prior where each component is uniformly distributed. (b)-(c) Informed posterior distributions for model *k* = 2,3, where the prior is taken to be the posterior distribution from model *k* − 1, as indicated by the arrows. Where appropriate, the posterior mode, or maximum likelihood estimate, is indicated with a red circle or sphere, and the boundary of the 95% credible region is indicated with a dashed red line. Modes for each model and univariate 95% credible intervals are given in Table 1. In all cases, the posterior density is scaled so that the maximum posterior density in the region shown is yellow, and blue represents a posterior density of zero.

With this information about *λ*, we now have two approaches to estimate *λ* and *D* from the circular barrier assay. First, we use the posterior in Figure 4(a) as a prior for *λ*, together with a uniform prior for *D*. This is the *informed* approach. The bivariate posterior in Figure 4(b) is well-defined, with little correlation between *D* and *λ*, and a mode of *D* = 620 *μ*m^2^/h and *λ* = 0.04 /h. Again, these estimates are consistent with previously-reported values (Treloar et al., 2013a; Haridas et al. 2017), but we note that previous studies have used extremely detailed experimental data that involves using nuclear stains to count individual cells and to construct detailed spatial and temporal distributions of cells within the circular barrier assay (Treloar et al., 2013a; Haridas et al. 2017). In contrast, here we simply use leading edge detection which completely circumvents the need for counting individual cells to construct detailed spatial and temporal distributions of cells in the barrier assay experiments. This means that our approach is very fast, simple-to-implement, and suitable for automation. In contrast, previous approaches are extremely labour intensive and cannot be easily automated (Treloar et al., 2013a; Haridas et al. 2017; Sengers et al., 2007; Cai et al., 2007; Vittadello et al. 2018).

In comparison with the informed approach, we now attempt to estimate *D* and *λ* directly with the leading edge data from the barrier assay with uniform priors for both *D* and *λ*. Indicative results in Figure 4(d) highlight several limitations with this approach. Here we have a very wide, poorly-defined posterior distribution with nonzero posterior density on the boundary of the parameter space. To arrive at this result we gradually widened the (*D, λ*) support, and it is important to note that the region in Figure 4(d), covers 0 < *λ* < 0.1 /h and 0 < *D* < 1000 *μ*m^2^/h. Since typical doubling times for cells are always in the range 10-20 h, it is clear that continuing to widen the support in Figure 4(d) will never lead to biologically-relevant parameter estimates. Therefore, we do not consider any further widening of the support. The reason that this approach fails to produce useful results is that the leading edge data alone is an insufficient summary statistic to identify *D* and *λ* from the barrier assay. Overall, comparing results in Figure 4(b) and Figure 4(d) confirm that our sequential Bayesian learning approach of combining minimal summary statistics from different experiments is both simple-to-implement and promising, as it leads to well-defined posterior distributions with a mode that is close to previously-determined estimates.

We now attempt to estimate *λ, D* and *δ* from the invasion assay. Again, with the informed approach, we use the posterior in Figure 4(b) as a prior for *λ* and *D*, with a uniform prior for *δ*. The posterior distribution in Figure 4(c) is well-defined, with a mode of *D* = 620 *μ*m^2^/h, *λ* = 0.04 /h and *δ* = 0.0036 /h. As before, these estimates for *D* and *λ* are consistent with previously-reported estimates, but we note that values of δ have not been reported previously for this kind of experimental data set. In contrast to the informed approach, result in Figure 4(e) show the outcome of using uniform priors for all three parameters, and we see that this leads to a poorly-defined posterior with regions of non-zero posterior density that are biologically irrelevant. We also reproduce the results from Figure 4 in the Supporting Material using synthetic experimental data to validate our approach. We examine a synthetic data set that uses a target parameter combination taken to be the maximum likelihood estimate from the informed posterior distribution for Model 3, and a synthetic data set where we lower the proliferation rate to simulate a more diffuse pattern of invasion. Full details on the method used to produce the synthetic experimental data sets are outlines in the Supporting Material. Similar trends are observed overall for both synthetic data sets. For example, the uninformed approach leads to poorly-defined posterior distributions, with regions of non-zero posterior density that are not close to the target parameter combination used to produce the synthetic data.

**Table 1.** Point estimates for each parameter, taken to be the posterior mode, or maximum likelihood estimate from the informed posterior distribution for each model. 95% credible intervals are estimated using the univariate marginal distributions and are shown in parentheses. All estimates are displayed to two significant figures.

| | $\lambda$ (/h) | $D$ ($\mu\text{m}^2/\text{h}$) | $\delta$ (/h) |
| --- | --- | --- | --- |
| Model 1 | 0.040 (0.038,0.042) | — | — |
| Model 2 | 0.040 (0.038,0.042) | 620 (480,800) | — |
| Model 3 | 0.040 (0.038,0.042) | 620 (480,800) | 0.0036 (0.0027,0.0046) |

Overall, comparing the informed posteriors in Figure 4(a)-(c) with the uninformed posteriors in Figure 4(d)-(e) we see the importance of the sequential approach. Given the full posterior distribution in Figure 4(c) we can integrate one of the components to form a series of three bivariate posterior distributions, as shown in Figure 5. Visually we see that *D* and *λ*, and *δ* and *λ* are approximately uncorrelated, whereas *δ* and *D* appear to be strongly negatively correlated. The Pearson correlation coefficients (Illowsky et al., 2015), given in Figure 5, confirm these visual observations.

**Fig. 5.**
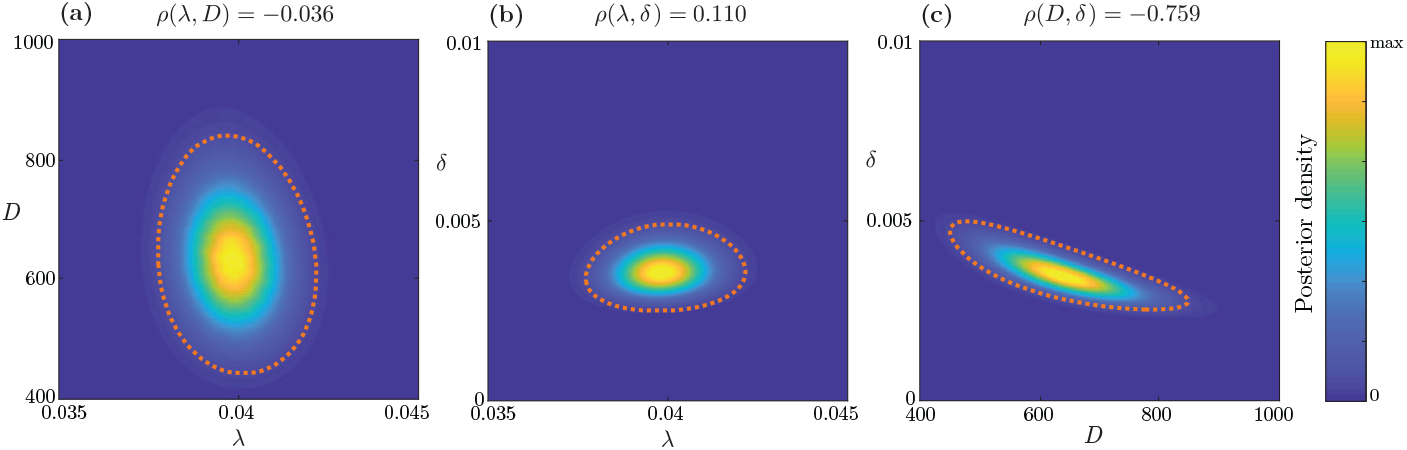
Informed bivariate marginal posterior distributions for model 3. Bivariate marginal posterior distributions formed by integrating out *δ, D* and *λ* in (a), (b) and (c), respectively, using the informed posterior distribution in Figure 4(e). The Pearson correlation coefficient, *ρ*(·, ·), is approximated using quadrature, and is shown, as indicated, for each marginal bivariate distribution. The 95% credible region for each bivariate marginal distribution is enclosed by the orange dashed lines.

In addition to visualising the posterior and marginal posterior distributions in Figure 4(c) and Figure 5, respectively, we can also calculate and compute the credible region shown in Figure 6.

**Fig. 6.**
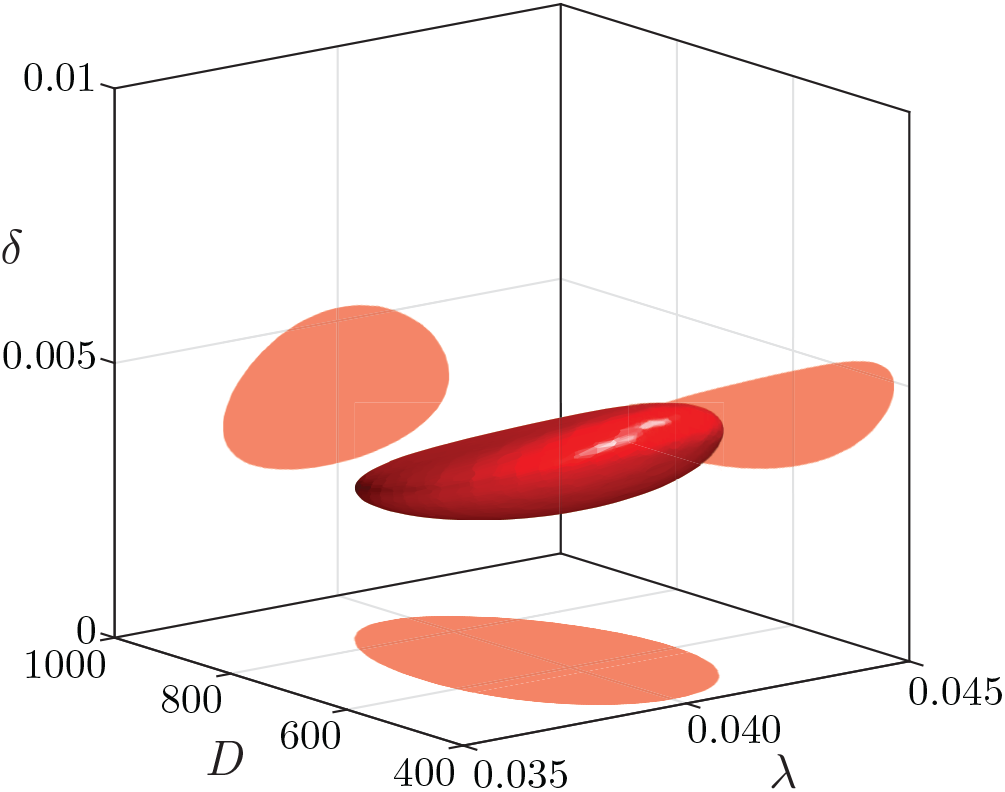
95% credible region for the model 3 posterior distribution. Given experimental data from all experiment types, we are 95% confident that the parameter combination for model 3 lies within this region. Note that shadows on axis planes do not indicate marginal posterior distributions, but rather the profile of the 95% credible region, viewed perpendicular to the plane. Estimates of the three bivariate marginal posterior distributions are given in Figure 5.

Now that we have arrived at a well-defined posterior distribution for ***Θ***, we can sample from this distribution and evaluate all three models and compare the expected model solution, and variability across many samples of model solutions, with the experimental data. A summary of the experimental data and model predictions, including 95% prediction intervals, are shown in Figure 7(a)-(c) for the proliferation assay, circular barrier assay and the invasion assay. For the proliferation assay, the barrier assay, and the invasion assay we see that the expected model predictions passes through most of the experimental data points. Again, for all three experiments we see that the 95% prediction intervals encompass almost all of the experimental measurements, as we would expect. In addition to showing how the solution of Equations (4)-(5) predicts the position of the leading edge of the invading population, min{*x*: *C*(*x, t*) = 0} in Figure 7(c), we also show the full solution of Equations (4)-(5) in Figure 7(d). Here we see the temporal evolution of both the melanoma density, *C*(*x, t*), and the density of skin tissues, *S*(*x, t*). These profiles show that the advance of the melanoma cell density profile in the positive *x* direction is closely associated with the retreat of the skin tissue profile, as expected. This coupling between the advance of the melanoma cells and the retreat of the skin tissues is evident in Figure 1(i)–(l). In the solution of the mathematical model, the simultaneous migration and proliferation of melanoma cells, coupled with the retreat of the tissue, gives rise to an advancing front of melanoma cells that is illustrated in the space-time diagram in Figure 7(e).

**Fig. 7.**
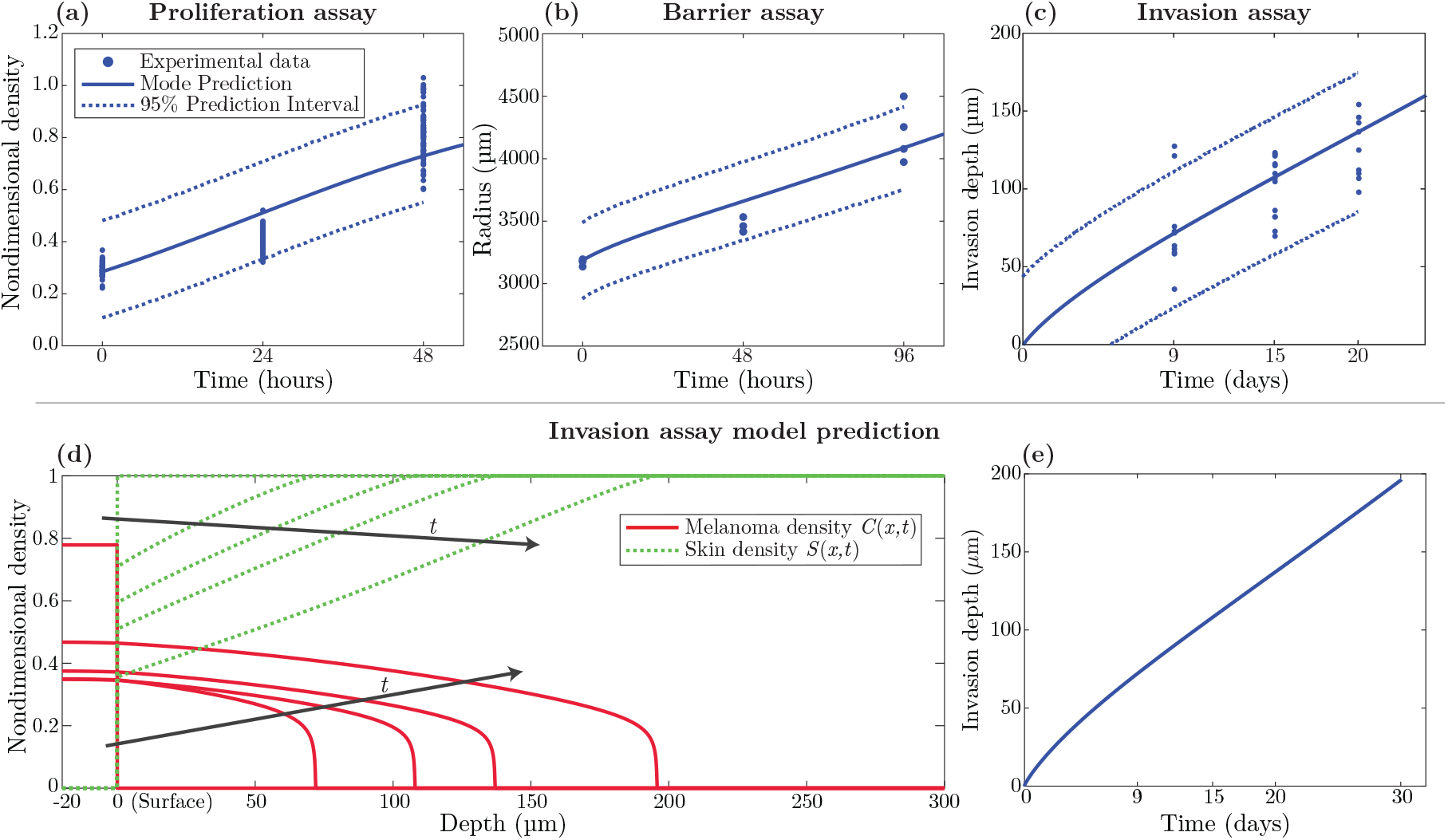
Experimental data and model predictions for all experiments. Experimental data and model predictions shown for experiment type 1, 2 and 3 in (a)-(c), respectively. Solid blue lines indicate model predictions using the mode parameter combination. Dashed blue lines in indicate approximate 95% prediction intervals. That is, the model predicts 95% of observations lie within this interval. (d) A detailed model prediction of the invasion assay using Equations (4)-(5) parameterised using the mode parameter combination. Density curves are shown as a function of depth for both simulated melanoma (red) and skin (blue) at times where experimental observations are taken, *t* = 9,15 and 20 days. Furthermore, an additional solution is shown at *t* = 30 days, which is beyond the timescale of the experimental data. The location of the melanoma cell front, which is the smallest depth such that *C*(*x,t*) = 0, is visible in (d) since the *C*(*x,t*) profile is sharp-fronted. The position of the front is shown as a function of time in (e).

## 6 Conclusion and Outlook

Continuum mathematical models of cell invasion typically involve coupled partial differential equations that describe how a population of cells degrades and simultaneously invades into some biological tissue. These models have become very well established in the literature over the last 20 years, and have been developed to describe malignant invasion (Fasano et al., 2009; Gatenby and Gawlinski, 1996; Marchant et al., 2001; Marchant and Norbury, 2002; Marchant et al., 2006; Perumpanani et al., 1999; Perumpanani et al., 2000; Smallbone et al., 2005) and invasion during embryonic development (Landman and Pettet, 1998; Landman et al., 2003; Landman et al., 2005; Simpson and Landman 2007). The scientific importance of these models (Gatenby and Gawlinski, 1996) and the mathematical analysis of these models (Perumpanani et al. 1999) are both relatively well advanced. Yet, it is surprising that despite the significance of these models, that there are presently no standard statistical protocols for calibrating these models using experimental data and/or experimental images. Our work is the first time these kinds of models have been quantitatively calibrated to match spatially quantitative measurements from invasion experiments.

In this work we present a Bayesian sequential learning approach, and demonstrate how it can be used to parameterise a simple model of cell invasion using data describing how a population of melanoma cells invades into human skin tissue. A key attraction of our approach is that we use images from a sequence of increasingly-sophisticated experiments. The measurements from each image are objective and straightforward, yet when these simple measures are combined sequentially, they allow us to parameterise the mathematical model to arrive at well-defined posterior distributions from which biologically-relevant parameter estimates can be taken. In contrast, taking a naive approach and simply estimating all parameters simultaneously from the images of the invasion assay leads to poorly-defined parameter estimates that, in this case, are biologically irrelevant.

While we have chosen to present our approach using a standard mathematical model of invasion in which we make fairly standard assumptions, it is possible to apply our approach to other types of models. For example, here we make the standard assumption that cells proliferate logistically in the invasion assay. However, if additional evidence suggested that a more general sigmoid growth model was appropriate (Browning et al., 2017; Sarapata et al., 2014), then our inference procedure could be applied to any other deterministic growth model. Similarly, we have used a non-linear diffusion term in the invasion model so that the diffusivity of the melanoma cells is a linearly decreasing function of total density. Again, if there were some evidence that some other kind of decreasing function of total density was warranted (Cai et al., 2007), our procedure could be repeated using a slightly different model with a different functional form for the nonlinear diffusivity.

Our inference approach is novel from a statistical point of view as we make progress by sequentially estimating parameters in a sequence of related models. This approach requires very little prior knowledge of the parameters and leads to well-defined posterior distributions. Deterministic models of cell migration and cell invasion are often calibrated to match experimental data using maximum likelihood, least-squares approaches (Cai et al., 2007; Sengers et al., 2007; Bowden et al., 2014; Hormuth et al., 2017). Such approaches produce a best-fit parameter combination but do not provide a means of systematically incorporating experimental variability from a sequence of related, but distinct experiments. As a result, parameter estimates produced using standard maximum likelihood approaches across a sequence of increasingly sophisticated models may not make sense. Our approach enforces a sensible relationship between those parameters estimated in the simpler experiments and those parameters estimated using more sophisticated experiments. The importance of taking a sequential approach is demonstrated in our study as we show that attempting to estimate all three parameters in the mathematical model using images from the invasion assay, without applying informed prior knowledge, leads to a poorly defined posterior distribution that may produce biologically irrelevant parameter combinations. Our informed approach may also be used in conjunction with gradient search techniques to find the maximum likelihood estimate. This could be advantageous in the sense that a gradient search technique does not require the computation of the posterior density over a wide range of the parameter space. However, the disadvantage of using such an approach is that it requires a good initial guess so that the initial guess lies relatively close to the maximum. The problem of having a good initial guess is particularly challenging because much of the posterior density in each of the distributions in 4 is essentially zero and extremely flat, meaning that gradient methods could suffer from convergence problems. Our approach avoids this issue because we simply expand the support of the search region in a computationally efficient way until we identify the maximum.

In our study, we focus on a likelihood-based technique as we are able to specify our likelihood function. Approximate techniques for parameter inference, such as approximate Bayesian computation (ABC), are also widely used to calibrate mathematical models to match experimental data (Toni et al., 2009; Beaumont et al., 2002; Browning et al., 2018), and are a necessity with stochastic mathematical models where the likelihood function is intractable. An extension of our study, that could include stochastic or individual based models, could apply such techniques such as ABC to our data set in a similar way. A key limitation of methods that rely on ABC is that they require a prior distribution to be fully specified before performing inference (Fearnhead et al., 2012). Another limitation is that ABC techniques typically require a large number of model realisations to produce a posterior distribution. While different variations of ABC have been developed to alleviate some of these limitations, such as Markov chain Monte Carlo sampling (Toni et al., 2009), sequential Monte Carlo sampling (Sisson et al., 2007), multilevel rejection sampling (Warne et al., 2018), and hierarchical ABC (Maclaren et al., 2017), our approach avoids some of these issues as we are able to directly specify the likelihood. For example, our application of a likelihood-based method enables the posterior density to be calculated numerically using a coarse discretisation of the parameter space using a relatively small number of deterministic model realisations. Such a coarse discretisation can be used to explore the posterior support, and the posterior distribution can be enhanced by refining the discretisation within the support, or by applying interpolation techniques. Finally, calculating the posterior distribution directly using a likelihood-based approach allows us to compute measures such as credible regions, and posterior mode estimates, without further data processing.

## Acknowledgements

This work is supported by the Australian Research Council (DP170100474). We appreciate helpful comments from David Warne, and two anonymous referees.

