## Supplementary Materials for "A Bayesian sequential learning framework to parameterise continuum models of melanoma invasion into human skin" (from another version of this preprint: version 1, 2018-03-19)

**Noname manuscript No.**
(will be inserted by the editor)

### Supplementary Material: A Bayesian sequential learning framework to parameterise continuum models of melanoma invasion into human skin

**Alexander P Browning** · **Parvathi Haridas** ·
**Matthew J Simpson**

Received: date / Accepted: date

Alexander P Browning
School of Mathematical Sciences, Queensland University of Technology (QUT), Brisbane, Australia

Parvathi Haridas
Institute of Health and Biomedical Innovation, QUT, Kelvin Grove, Australia

Matthew J Simpson
School of Mathematical Sciences, Queensland University of Technology (QUT), Brisbane, Australia


#### Mathematical model and simplification

Here we present a typical continuum model describing just the key features though to be relevant to cellular invasion in the context of melanoma invading human skin. In particular, we are inspired by the model presented by Perumpanani et al. (1999) who consider a one-dimensional, three-species system where the dependent variables represent:

1. a population of cells with density, $c(x,t)$;
2. a substrate with density $s(x,t)$; and
3. a concentration of protease, $p(x,t)$.

For our work we interpret $c(x,t)$ as the density of melanoma cells, $s(x,t)$ as the density of de-epidermised human skin tissue, and $p(x,t)$ representing a chemical, such as protease, produced by the melanoma cells that acts to degrade the de-epidermised tissue.

To formulate a mathematical model we make the following minimal assumptions:

1. melanoma cells undergo random, diffusive migration;
2. melanoma cells proliferate;
3. contact inhibition of migration is important, and is modelled using a nonlinear diffusivity function so that the net melanoma cell diffusivity is a linearly decreasing function of the substrate density;
4. contact inhibition of proliferation is relevant, and modelled so that the net proliferation rate of melanoma cells is a linearly decreasing function of total cellular and substrate density so that the melanoma cells are immobile when the total density is at the carrying capacity density;
5. consistent with previous continuum models of cellular invasion (Perumpanani et al., 1999), we assume there is no net flux of the substrate, and that the substrate decays in the presence of the protease; and

6. there is no net flux of the protease, that the protease is produced by melanoma cells, and that the protease undergoes linear decay (Perumpanani et al., 1999).

Combining these assumptions, and supposing that the natural maximum density of the melanoma cells plus the substrate is given by some carrying capacity density, $k > 0$, in one-dimensional Cartesian coordinates we obtain,

$$\frac{\partial c(x,t)}{\partial t} = D\frac{\partial}{\partial x}\left[\left(1-\frac{s(x,t)}{k}\right)\frac{\partial c(x,t)}{\partial x}\right] + \lambda c(x,t)\left[\left(1-\frac{c(x,t)+s(x,t)}{k}\right)\right], \quad (1)$$

$$\frac{\partial s(x,t)}{\partial t} = -ls(x,t)p(x,t), \quad (2)$$

$$\frac{\partial p(x,t)}{\partial t} = mc(x,t) - np(x,t), \quad (3)$$

where $D$ is the low-density diffusivity of melanoma cells, $\lambda$ is the low-density proliferation rate of melanoma cells, $l$ is the rate at which the substrate decays per unit of protease, $m$ is the rate at which melanoma cells produce protease, and $n$ is the natural decay rate of the protease. Despite the many simplifications invoked to arrive at Equations (1)-(3), to use this model to quantitatively match and predict experimental observations we must deal with three dependent variables ($c(x,t)$, $s(x,t)$ and $p(x,t)$), and six free parameters ($D$, $\lambda$, $k$, $l$, $m$ and $n$). Following Perumpanani et al., (1999) and many others (e.g. Breward et al., 2002), we simplify the model by assuming that the timescale of protease degradation is the fastest process in the system. Invoking a quasi-steady assumption by setting $\partial p(x,t)/\partial t = 0$, we obtain $p(x,t) = mc(x,t)/n$. Under this assumption, the number of dependent variables reduces to two, and the number of free parameters to three,

$$\frac{\partial c(x,t)}{\partial t} = D\frac{\partial}{\partial x}\left[\left(1-\frac{s(x,t)}{k}\right)\frac{\partial c(x,t)}{\partial x}\right] + \lambda c(x,t)\left[\left(1-\frac{c(x,t)+s(x,t)}{k}\right)\right], \quad (4)$$

$$\frac{\partial s(x,t)}{\partial t} = -\gamma s(x,t)c(x,t), \quad (5)$$

where $\gamma = lm/n$. To further simplify the model we non-dimensionalise the dependent variables by considering a non-dimensional cell density, $C(x,t) = c(x,t)/k$, and a non-dimensional substrate density, $S(x,t) = s(x,t)/k$, giving

$$\frac{\partial C(x,t)}{\partial t} = D\frac{\partial}{\partial x}\left[(1-S(x,t))\frac{\partial C(x,t)}{\partial x}\right] + \lambda C(x,t)\left[1-C(x,t)-S(x,t)\right], \quad (6)$$

$$\frac{\partial S(x,t)}{\partial t} = -\delta C(x,t)S(x,t), \quad (7)$$

where, $\delta = \gamma k$. In this format, the model involves two non-dimensional dependent variables ($C(x,t)$ and $S(x,t)$), and three free parameters ($D$, $\lambda$ and $\delta$).

We freely acknowledge that this mathematical model involves several assumptions, and that each of these assumptions could be revisited in future studies if there were experimental evidence to suggest that such a revision is warranted. For example, here we consider that the net proliferation rate of melanoma cells is a linearly decreasing function of the total density. Other studies suggest that some kind of non-linear decreasing function could lead to an improved match with experimental data (Browning et al., 2017; Sarapata et al., 2014). However, here we are focusing on using a mathematical model that involves just the essential biological features that are though to be relevant to the system of interest, and we explain how to obtain parameter estimates for this model. This question is important because we are unaware of any previous study that provides quantitative parameter estimate for this kind of model using experimental measurements and observations. Importantly, as we show, there is sufficient redundancy in the present model that it is challenging to identify unique combinations of $D$, $\lambda$ and $\delta$ without great care. This challenge would be made more difficult if we chose to work with a more general model that contained additional exponents that also require estimation (Browning et al., 2017; Sarapata et al., 2014). Therefore, we leave the question of model selection for future consideration.

##### Hierarchy of mathematical models

Embedded within Equations (6)–(7) is a hierarchy of models:

Model 3: spatial and temporal two-component model

The full two-component system given by,

$$\frac{\partial C(x,t)}{\partial t} = D\frac{\partial}{\partial x}\left[(1-S(x,t))\,\frac{\partial C(x,t)}{\partial x}\right] + \lambda C(x,t)\,[1-C(x,t)-S(x,t)]\,, \quad (8)$$

$$\frac{\partial S(x,t)}{\partial t} = -\delta C(x,t)S(x,t). \quad (9)$$

Model 2: spatial and temporal one-component model

If we consider a population of cancer cells in the absence of the substrate by setting $S(x,t) = 0$, we obtain

$$\frac{\partial C(x,t)}{\partial t} = D\frac{\partial^2 C(x,t)}{\partial x^2} + \lambda C(x,t)\,[1-C(x,t)]\,, \quad (10)$$

which is the celebrated Fisher-Kolmogorov equation. To apply the model to match our experimental observations, we make use of the fact that we have circular barrier assays in which the geometry is two-dimensional with radial symmetry. Therefore, it is appropriate to re-cast the model in a radial co-ordinate system, given by

$$\frac{\partial C(r,t)}{\partial t} = \frac{D}{r}\frac{\partial}{\partial r}\left[r\frac{\partial C(r,t)}{\partial r}\right] + \lambda C(r,t)\,[1-C(r,t)]\,, \quad (11)$$

where $r$ is the radial coordinate.

Model 1: temporal one-component model

Again, for a population of cancer cells in the absence of the substrate we set $S(x,t) = 0$. Furthermore, if we consider a spatially uniform population of cancer cells we have $\partial C/\partial x = 0$, and so $C(x,t)$ can be written as $C(t)$, giving

$$\frac{\mathrm{d}C(t)}{\mathrm{d}t} = \lambda C(t)\,[1-C(t)]\,, \quad (12)$$

which is the well-known logistic equation, which has the solution

$$C(t) = \frac{C(0)}{C(0) + \mathrm{e}^{-\lambda t}(1 - C(0))}. \tag{13}$$

Here we have listed these models in a decreasing order of mathematical and biological complexity. This ordering makes it clear that Model 1 is a direct simplification of Model 2, and that Model 2 is a direct simplification of Model 3.

#### Numerical methods

We solve models 2 3 using a finite difference method with constant grid spacing, and implicit Euler time stepping with constant time steps of duration, $\delta t$. For model 3, in Cartesian coordinates, a uniform discretization is used where node spacing is $\delta x$. For model 2, in radial coordinates, a uniform discretization is used where the node spacing is $\delta r$. Picard iteration with convergence tolerance $\varepsilon$ is used to solve the resulting systems of nonlinear equations. For all results presented here, we are careful to choose $\delta x$, $\delta r$, $\delta t$ and $\varepsilon$ to be sufficiently small so that the numerical solutions are grid-independent.

##### Numerical discretisation of model 2

Here, we consider the domain, $0 < r < R$ into $N$ equally-sized intervals of width $\delta r$, where $R$ is sufficiently large so that $C(R,t) = 0$ for $t > 0$. We denote the density, $C(r_i,t)$ at the $i$th node as $C_i^{(t)}$, such that $r_i = i\delta r$. Applying a backward Euler discretisation to Equation (11) we obtain

$$\begin{aligned} \frac{C_i^{(t+\delta t)} - C_i^{(t)}}{\delta t} = \frac{D}{r_i \delta r}\Bigg\{ & r_{i+\frac{1}{2}} \left[ \frac{C_{i+1}^{(t+\delta t)} - C_i^{(t+\delta t)}}{\Delta r} \right] \\ & - r_{i-\frac{1}{2}} \left[ \frac{C_i^{(t+\delta t)} - C_{i-1}^{(t+\delta t)}}{\Delta r} \right] \Bigg\} + \lambda C_i^{(t+\delta t)} (1 - C_i^{(t+\delta t)}), \end{aligned} \tag{14}$$

for the internal nodes, that is, $1 < i < N$. We apply no-flux boundary conditions by enforcing

$$\begin{aligned} C_i^{(t)} &= C_{i+1}^{(t)},\ i = 0, \\ C_i^{(t)} &= C_{i-1}^{(t)},\ i = N+1. \end{aligned}$$

Writing Equation (14) as

$$C_i^{(t)} = \alpha_i C_{i-1}^{(t+\delta t)} + \beta_i C_i^{(t+\delta t)} + \gamma_i C_{i+1}^{(t+\delta t)},$$

we have,

$$\begin{aligned} \alpha_i &= \frac{D\Delta t}{2r_i\delta r} - \frac{D\delta t}{\delta r^2}, \\ \beta_i &= 1 + 2\frac{D\delta t}{\delta r^2} + \lambda\delta t(1 - C_i^{(t+\delta t)}), \\ \gamma_i &= -\frac{D\delta t}{2r_i\delta r} - \frac{D\delta t}{\delta r^2}, \end{aligned}$$

for all internal nodes, and $\beta_0 = \beta_N = 1$, $\gamma_0 = \alpha_N = -1$ which enforces the appropriate boundary conditions. We may then write Equation (14) in matrix form

$$\mathbf{A}(\mathbf{C}^{(t+\delta t)})\mathbf{C}^{(t+\delta t)} = \mathbf{C}^{(t)}, \tag{15}$$

where $\mathbf{C}^{(t)} = \langle C_0^{(t)}, C_1^{(t)}, ..., C_N^{(t)} \rangle$ and $\mathbf{A}(\mathbf{C}^{(t+\delta t)})$ is a tridiagonal matrix where $\alpha_i, \beta_i$ and $\gamma_i$ form the lower diagonal, main diagonal and upper diagonal. We solve Equation (22) using Picard iteration.

For model 2, we set $\delta r = 1$ $\mu$m, $\delta t = 0.1$ /h and $\varepsilon = 10^{-5}$. This choice of discretisation leads to grid-independent results for the parameter values that we consider.

Numerical discretisation of model 3

Similar to our discretisation of model 2, we divide the domain, $0 < x < L$ into $N$ equally-sized subintervals of width $\delta x$, where $L$ is sufficiently large such that $C(L,t) = 0$ for $t > 0$. We denote the density of melanoma cells, $C(x_i,t)$ at the $i$th node, as $C_i^{(t)}$, and the density of skin tissue, $S(x_i,t)$ at the $i$rh node, as $S_i^{(t)}$ such that $x_i = i\delta x$. Applying a backward Euler discretisation to Equation (8) and Equation (9) we obtain

$$\begin{aligned}\frac{C_i^{(t+\delta t)} - C_i^{(t)}}{\delta t} = \frac{D}{2\delta x^2}\Bigg\{ & \left(f(S_i^{(t+\delta t)}) + f(S_{i+1}^{(t+\delta t)})\right)\left(C_{i+1}^{(t+\delta t)} - C_i^{(t+\delta t)}\right) \\ & - \left(f(S_i^{(t+\delta t)}) + f(S_{i-1}^{(t+\delta t)})\right)\left(C_i^{(t+\delta t)} - C_{i-1}^{(t+\delta t)}\right)\Bigg\} \\ & +\lambda C_i^{(t+\delta t)} g(C_i^{(t+\delta t)}, S_i^{(t+\delta t)})),\end{aligned} \tag{16}$$

and,

$$\frac{S_i^{(t+\delta t)} - S_i^{(t)}}{\delta t} = \delta S_i^{(t+\delta t)} C_i^{(t+\delta t)} \tag{17}$$

where $f(S_i) = 1 - S_i$, $g(C_i, S_i) = 1 - C_i - S_i$ and for the internal nodes, $1 < i < N$. We apply no-flux boundary conditions by enforcing

$$\begin{aligned} C_i^{(t+\delta t)} &= C_{i+1}^{(t+\delta t)},\ i = 0, \\ C_i^{(t+\delta t)} &= C_{i-1}^{(t+\delta t)},\ i = N+1. \end{aligned}$$

Writing Equation (16) as

$$C_i^{(t)} = \alpha_i C_{i-1}^{(t+\delta t)} + \beta_i C_i^{(t+\delta t)} + \gamma_i C_{i+1}^{(t+\delta t)}, \tag{18}$$

we obtain

$$\alpha_i = -\frac{D\delta t}{2\delta x^2}\left(f(S_{i-1}^{(t+\delta t)}) + f(S_i^{(t+\delta t)})\right) \tag{19}$$

$$\beta_i = 1 + \frac{D\delta t}{2\delta x^2}\left(f(S_{i-1}^{(t+\delta t)}) + 2f(S_i^{(t+\delta t)}) + f(S_{i+1}^{(t+\delta t)})\right) - \lambda\,\delta t g(C_i^{(t+\delta t)}, S_i^{(t+\delta t)})) \tag{20}$$

$$\gamma_i = -\frac{D\delta t}{2\delta x^2}\left(f(S_i^{(t+\delta t)}) + f(S_{i+1}^{(t+\delta t)})\right) \tag{21}$$

on the internal nodes, and $\beta_0 = \beta_N = 1$, $\alpha_0 = \alpha_N = -1$ which enforces the boundary conditions. We may then write Equation (16) as a matrix system

$$\mathbf{A}(\mathbf{C}^{(t+\delta t)}, \mathbf{S}^{(t+\delta t)})\mathbf{C}^{(t+\delta t)} = \mathbf{C}^{(t)}, \tag{22}$$

where $\mathbf{C}^{(t)} = \langle C_0^{(t)}, C_1^{(t)}, ..., C_N^{(t)} \rangle$, $\mathbf{S}^{(t)} = \langle S_0^{(t)}, S_1^{(t)}, ..., S_N^{(t)} \rangle$ and $\mathbf{A}$ is a tridiagonal matrix where $\alpha_i, \beta_i$ and $\gamma_i$ form the lower diagonal, main diagonal and upper diagonal. We may then solve Equation (17) and Equation (22) sequentially using Picard iteration.

For model 3, we use $\delta x = 0.05$ $\mu$m, $\delta t = 0.01$ days and $\varepsilon = 10^{-5}$. This choice of discretisation leads to grid-independent results for the parameter values that we consider.

#### Enhancing resolution of posterior distributions using interpolation

Since models 2 and 3 require numerical algorithms to solve the partial differential equations, we calculate the posterior density on a relatively coarse square discretisation of the parameter space. The discretisation for each model and each parameter are summarised in Table 1. These choices of discretisations are made such that we obtain a point estimate of each parameter to two significant figures of accuracy.

All posterior distributions we deal with appear to be smooth. Therefore, we refine our estimates of the posterior distributions using a spline interpolation (Mathworks,

2018). To achieve this we discretise the parameter space using 500 equally-spaced points in each dimension. To illustrate this procedure, Figure 1a shows a relatively coarse, posterior distribution for model 2, with a resolution of 35 $\times$ 61 estimates of the posterior density in the $(D, \lambda)$ parameter space. Visually, this coarse discretisation appears to be reasonably well behaved as we appear to have covered the support of the $(D, \lambda)$ parameter space and the distribution of non-zero posterior density appears to be reasonably smooth. Figure 1b shows the associated enhanced posterior distribution that is used to compute estimate the credible regions.

| | Model 1 | Model 2 | | Model 3 | |
|---|---|---|---|---|---|
| Parameter | Uninformed | Uninformed | Informed | Uninformed | Informed |
| $\lambda$ | $1\times10^{-5}$ | $5\times10^{-4}$ | $5\times10^{-5}$ | $3\times10^{-3}$ | $3\times10^{-4}$ |
| $D$ | – | 5 | 5 | 20 | 10 |
| $\delta$ | – | – | – | $3\times10^{-3}$ | $1\times10^{-4}$ |

**Table 1** Discretisations of the parameter space.

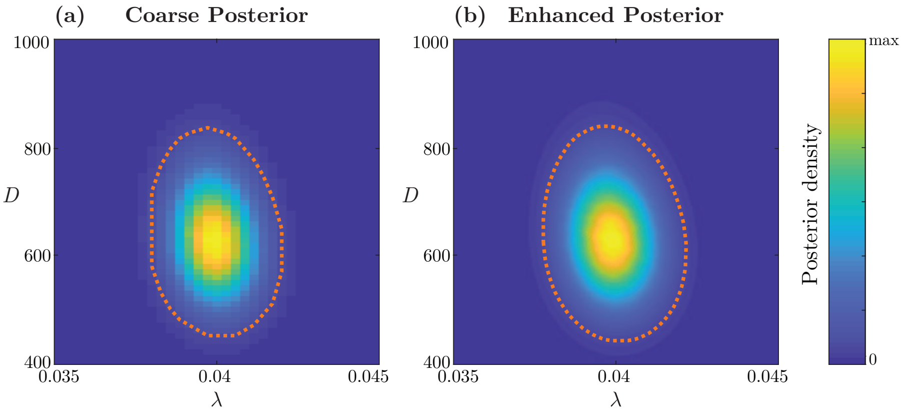


**Fig. 1 Enhancing the resolution of posterior distributions.** (a) Original, coarse posterior distribution with a resolution of 35 $\times$ 61. (b) Enhanced posterior distribution. `interp` has been used to increase the resolution to 500 $\times$ 500 by calculating posterior densities at intermediate grid points using a spline interpolation (Mathworks, 2018). The 95% credible region for each bivariate marginal distribution is enclosed by the orange dashed lines.
